# Inducing representational change in the hippocampus through real-time neurofeedback

**DOI:** 10.1101/2023.12.01.569487

**Authors:** Kailong Peng, Jeffrey D. Wammes, Alex Nguyen, Marius Cătălin Iordan, Kenneth A. Norman, Nicholas B. Turk-Browne

**Affiliations:** Department of Psychology, Interdepartmental Neuroscience Program, Yale University; Department of Psychology, Centre for Neuroscience Studies, Queen’s University; Department of Psychology, Princeton Neuroscience Institute, Princeton University; Department of Brain and Cognitive Sciences, Department of Neuroscience, University of Rochester; Department of Psychology, Princeton Neuroscience, Institute Princeton University; Department of Psychology, Wu Tsai Institute, Yale University

**Keywords:** Real-time fMRI, Nonmonotonic plasticity, Closed-loop neurofeedback, High-performance computing, Psychophysics, Machine learning

## Abstract

When you perceive or remember one thing, other related things come to mind. This competition has consequences for how these items are later perceived, attended, or remembered. Such behavioral consequences result from changes in how much the neural representations of the items overlap, especially in the hippocampus. These changes can reflect increased (integration) or decreased (differentiation) overlap; previous studies have posited that the amount of coactivation between competing representations in cortex determines which will occur: high coactivation leads to hippocampal integration, medium coactivation leads to differentiation, and low coactivation is inert. However, those studies used indirect proxies for coactivation, by manipulating stimulus similarity or task demands. Here we induce coactivation of competing memories in visual cortex more directly using closed-loop neurofeedback from real-time fMRI. While viewing one object, participants were rewarded for implicitly activating the representation of another object as strongly as possible. Across multiple real-time fMRI training sessions, they succeeded in using the neurofeedback to induce coactivation. Compared with untrained objects, this coactivation led to behavioral and neural integration: The trained objects became harder for participants to discriminate in a categorical perception task and harder to decode from patterns of fMRI activity in the hippocampus.

## Introduction

Unlike a hard drive that stores files in dedicated blocks, the brain partially re-uses neurons that already store existing memories when forming a new memory. Because of the resulting overlap, when we later attempt to retrieve one memory, other related memories can come to mind. The competition between these coactivated memories can cause interference and behavioral errors in the moment. At the same time, such coactivation can also induce learning, with consequences for how the target and competitor memories are subsequently represented [1, 2, 3]. Namely, this learning can increase the amount of subsequent overlap between the neural populations representing each memory (integration) or it can reduce overlap (differentiation).

Both integration and differentiation have been reported at the level of distributed representations captured by functional magnetic resonance imaging (fMRI). Integration has been observed in statistical learning tasks in which arbitrarily chosen objects reliably co-occur in time or space [4, 5, 6, 7]. For example, when paticipants view a continuous sequence of fractal images containing temporal pairs, in which one image is always followed by another, the paired images come to evoke more similar patterns of fMRI activity in the hippocampus [4]. Integration has also been observed in tasks involving deliberate associative learning: Linking two arbitrary cues to a common associate increases the similarity of fMRI activity patterns for the two cues in the hippocampus and other regions [8, 9, 10, 11]. Strikingly, other, closely related studies (and, in some cases, closely related conditions within the same studies) have observed the exact opposite result of differentiation [12, 13, 11, 14, 15, 16, 4, 17]. For example, multiple studies have found that pairing two cues with the same associate can lead to differentiation in the hippocampus [15, 11].

Reconciling these discrepant findings poses a major theoretical challenge. According to a Hebbian learning process, coactivation between memories should strengthen synaptic connections to and among the neural populations these memories share, resulting in integration that scales monotonically with the degree of coactivation. However, this cannot explain why pairing two scenes with the same face (which acts as a “bridge”, allowing activation to spread between the scenes) results in lower hippocampal similarity than when the scenes are paired with different faces and thus lack the bridge [15]. This is one of several cases where manipulations that should increase coactivation have resulted in decreased neural similarity [18, 4, 16, 14, 12, 11, 17, 19]. These kinds of results can potentially be explained by the non-monotonic plasticity hypothesis (NMPH), which posits that representational change follows a U-shaped pattern as a function of memory coactivation [3]. Under this hypothesis, the baseline case of no coactivation leads to no representational change, moderate coactivation leads to subsequent differentiation, and high coactivation leads to integration. A computational model that implements this learning rule was able to reproduce patterns of behavioral and neural change from three recent studies [20].

Although several studies have reported findings consistent with NMPH, only one study has attempted to parametrically manipulate the amount of coactivation to trace the predicted U-shaped curve [14]. In this study, a convolutional neural network was used to synthesize eight pairs of images that varied parametrically in visual similarity. The similarity structure of these images was validated by showing that fMRI pattern similarity in the visual cortex scaled with increasing model similarity. These pairs were then embedded in a statistical learning task, with the logic that overlap between pairmates in their cortical inputs to the hippocampus would drive coactivation that (in turn) would have a non-monotonic relationship to how the representations would change. Namely, low visual similarity would lead to weak coactivation in the hippocampus and no representational change, medium similarity would lead to moderate coactivation and differentiation, and high similarity would lead to strong coactivation and integration. The fMRI pattern of activity evoked by each image in the hippocampus was measured before and after statistical learning. The dentate gyrus (DG) subfield in particular exhibited a non-monotonic result with clear differentiation for pairs with medium (but not low or high) visual similarity. The role of DG is consistent with prior studies [13, 17] and with computational models suggesting that DG and CA3 may be biased to differentiation whereas CA1 is biased to integration [21]. Indeed, integration was not obtained for pairs with high visual similarity, suggesting that these images may not have triggered strong enough coactivation.

The current study builds on this work by providing a more direct test of how coactivation drives representational change in the hippocampus. Rather than relying on stimulus manipulations or other indirect proxies to induce coactivation in regions of the visual cortex providing hippocampal input, here we manipulate coactivation itself with closed-loop neurofeedback. This approach is grounded in impressive developments in the field of real-time fMRI over the past decade [22]. Real-time fMRI refers to conducting an fMRI study in which the blood oxygenation level-dependent (BOLD) data are analyzed immediately after collection, such that the resultant estimates of brain activity can be provided as feedback to the participant or used to modify the task in a closed-loop manner [23, 24]. This technique has found increasing success in clinical studies, to modulate disordered brain activity such as in depression [25, 26], and for basic science discovery, to train or test theories of cognitive abilities such as sustained attention [27], perceptual learning [28], category learning [29], and episodic memory [30].

Here we use real-time fMRI to induce coactivation of two objects in the visual cortex across three neurofeedback sessions, and we test how this impacts the similarity of their representations in the hippocampus in two additional sessions, before and after neurofeedback, respectively. To measure representational change in these pre and post-sessions, we take snapshots of how well the objects can be discriminated in behavior (using a categorical perception task) and in the brain (using multivariate pattern similarity). During neurofeedback sessions, coactivation is achieved by training the participant to incept activation of one “competitor” object (e.g., a chair) while another “presented” object (e.g., bed) is shown on the screen. To the extent that participants learn to strongly activate the competitor, we hypothesize that the representations of the presented and competitor objects will integrate, leading to a shallower discrimination slope in the behavioral categorical perception task (i.e., more confusion) and greater pattern similarity in the hippocampus, perhaps especially in the CA1 subfield, when comparing post to pre snapshots. If instead participants achieve only moderate competitor activation through neurofeedback, NMPH predicts that the representations of the presented and competitor objects will differentiate, leading to a steeper behavioral discrimination slope (i.e., easier to tell apart) and reduced hippocampal pattern similarity, possibly in DG and CA3 subfields. Beyond assessing change over time from pre to post-learning, we also include an additional baseline condition of two other objects (e.g., a table and a bench) for which we obtain behavioral and neural snapshots in the same way as the presented and competitor objects; crucially, these baseline objects are never subject to real-time fMRI training of coactivation, so we expect neither integration nor differentiation to occur in the hippocampus for these objects.

## Methods

### Participants

A total of 20 healthy young adults (mean age = 23.55; age range = 19–28; 14 female) with normal or corrected-to-normal visual acuity participated in this study. They completed five fMRI sessions lasting 1.5 hours each and were paid $20/hour. Sessions were scheduled on separate days but as close together as possible, from a minimum of five days in a row to a total span of eight days.

### Ethics Statement

All participants provided informed consent to a protocol approved by the Institutional Review Board at Yale University.

### Data acquisition

Data were acquired using a 3T Siemens Prisma scanner with a 64-channel head coil at the Brain Imaging Center at Yale University. For recognition and feedback functional runs, an echo-planar imaging (EPI) sequence was used to collect BOLD data (TR=2 s; TE=30 ms; voxel size=3 mm isotropic; FA=90°; IPAT GRAPPA acceleration factor=2; distance factor=25%), yielding 36 axial slices. Each recognition run contained 145 volumes and each feedback run contained 176 volumes. Two field map volumes (TR=5 s; TE=80 ms; otherwise matching the EPI scans) were acquired in opposite phase encoding directions. For anatomical alignment and visualization, we collected a 3D T1-weighted magnetization-prepared rapid acquisition gradient echo (MPRAGE) scan (TR=2.5 s; TE=2.9 ms; voxel size=1 mm isotropic; FA=8°; 176 sagittal slices; IPAT GRAPPA acceleration factor=2), and a 3D T2-weighted fast spin echo scan with variable flip angle (TR=3.2 s; TE=565 ms; voxel size=1 mm isotropic; 176 sagittal slices; IPAT GRAPPA acceleration factor=2).

### Real-time system

After image reconstruction, the DICOM files were streamed in real-time to Milgram, a high-performance cluster mounted on the Siemens console. The RT-Cloud software package [31] was used for preprocessing and analysis of each image, with the results transmitted to the task computer at the scanner over the network. This output was used to update the task on the next time point.

### Data preprocessing

For real-time analyses, each new DICOM file was aligned with 3dvolreg [32] to a template volume acquired from the middle volume of the first recognition run in the current session. The BOLD activity of every voxel was subjected to z-scoring over time using the running mean and standard deviation from the prior TRs in the current run. The data were masked to include only voxels in the region of interest (ROI) used for feedback prior to classifier training and testing. For offline analyses, the functional data were field map corrected with the topup tool in FSL [33], registered to the middle volume of the current run with MCFLIRT [34], and z-scored based on the full-time series from that run.

### Study design

The study consisted of five sessions (Table 1). Session 1 contained eight recognition runs in which the presented, competitor, and control objects were shown repeatedly. These data were used to train classifier models that could distinguish between all pairs of objects. The trained models were then tested on the feedback runs in Sessions 2, 3, and 4 to measure the amount of evidence for the competitor object during the viewing of the presented object. Participants were encouraged to increase the activation level of the competitor through adaptive, closed-loop neurofeedback, inducing coactivation between the presented and competitor objects. Session 5 mirrored the first session with eight recognition runs, in order to assess representational change.

**Table 1:** Multi-session study protocol for each participant.

| Day | Session 1 (~1.5h) | Session 2 (~1.5h) | Session 3 (~1.5h) | Session 4 (~1.5h) | Session 5 (~1.5h) |
| --- | --- | --- | --- | --- | --- |
| Task order | 8 recognition runs | 2 recognition runs | 2 recognition runs | 2 recognition runs | 8 recognition runs |
|  |  | 10 feedback runs | 10 feedback runs | 10 feedback runs |  |
|  |  | 2 recognition runs | 2 recognition runs | 2 recognition runs |  |

The assignment of objects to conditions was counterbalanced across participants in two batches. For the 10 participants in Batch 1, the presented object was bed, the competitor object was chair, and the control objects were table and bench (Fig. 1a). For the remaining 10 participants in Batch 2, the assignments were reversed with table as presented object, bench as the competitor object, and bed and chair as control objects.

**Figure 1:**
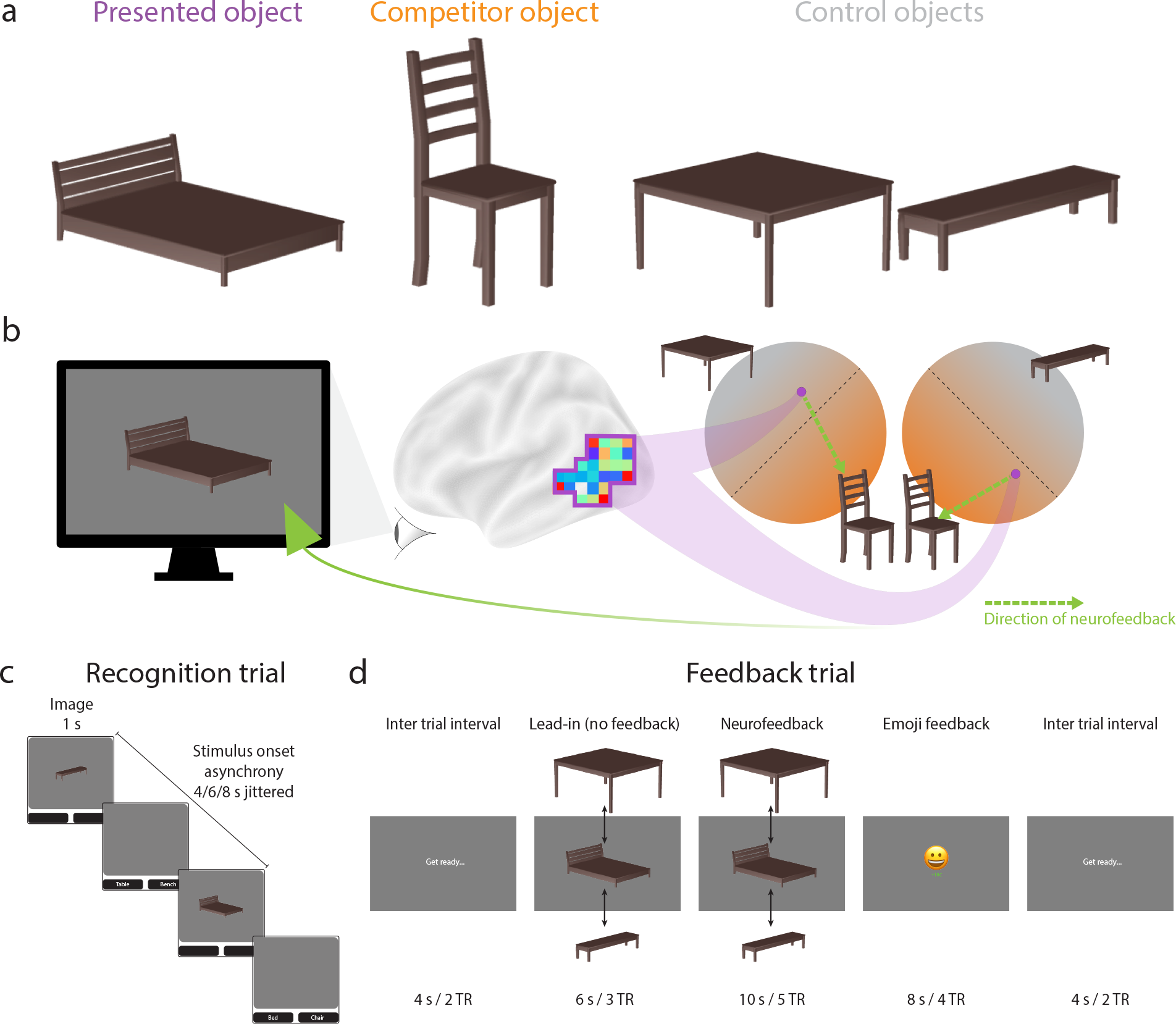
Study design. (a) Each participant received two objects for neurofeedback and the other two objects served as a baseline. (b) The presented object (e.g., bed) was shown on neurofeedback trials and began oscillating in size and shape. The goal of the participant was to make this wobbling stop, which they could achieve by activating the representation of the competitor object (e.g., chair) in their mind. Evidence for the competitor object was quantified based on a classifier trained to decode the competitor object relative to two control objects (e.g., table, bench). The amount of classifier evidence for the competitor needed to reduce the magnitude of wobbling was staircased to maximize coactivation between these objects. (c) Recognition trials showed one of the four objects at a time with no neurofeedback. These trials were used to train the classifier models and to measure neural snapshots of how the object is presented in the hippocampus. (d) Feedback trials were collected during real-time fMRI in order to induce coactivation. The main neurofeedback occurred during the object presentation (the amount of wobbling), though participants also received a valenced emoji based on their performance relative to their staircase.

### Recognition runs

During each trial of the recognition runs (Fig. 1c), participants were presented with one of four rendered furniture objects (bed, bench, chair, table) in one of several potential viewing angles [35, 36]. After 1 s, the object was removed from the screen and two furniture labels appeared below, from which participants had to choose which matched the object with an MRI-compatible button box. This response occurred during a jittered interval between trials of 4, 6, or 8 s. Each recognition run contained 48 trials, allowing for 12 repetitions per object and run. Three repetitions of each object appeared in each quarter of a run (no back-to-back repetitions), to ensure an even distribution of objects over time.

The data from the recognition runs in Session 1 were used to train six binary classifiers, corresponding to all combinations of the four objects. We used logistic regression classifiers with L2-norm regularization (penalty=1). Each of the six classifiers contrasted one pair of objects (e.g., chair vs. bench). The training data consisted of patterns of BOLD activity across voxels in a feedback ROI (described below) extracted 4 s after the onset of the object on each trial to account for the hemodynamic lag, labeled by the identity of the object.

### Feedback runs

During each trial of the feedback runs (Fig. 1d), the presented object was shown dynamically on the screen, appearing to wobble in size and shape [29]. BOLD activity patterns were extracted from the feedback ROI and supplied as input to the classifiers. To determine the activation level of the competitor object (Fig. 1b), we averaged the output of the two classifiers trained to discriminate the competitor object from each of the control objects (i.e., competitor vs. control1; competitor vs. control2). Similarly, the activation level of the presented object was determined by averaging the output of the two classifiers trained to discriminate the presented object from each of the control objects (i.e., presented vs. control1; presented vs. control2). Note that we did not consider the output of the classifier directly pitting presented vs. competitor objects because we wanted separate estimates of the evidence for these objects. For example, if both were active, the classifier output may be at chance, but this result is also possible if neither is active. Instead, we rely on the control objects to provide a neutral baseline for both presented and competitor objects.

Participants received multiple forms of feedback to help motivate them to increase the activation level of the competitor object and foster its coactivation with the presented object, including visual feedback via wobbling, monetary feedback via an increase in their bonus compensation, and valenced feedback via an emoji. If participants successfully raised the activation level of the competitor above an adaptive threshold, the magnitude of wobbling decreased: wobbling began each feedback trial (consisting of 5 TRs) at level 13 (maximal), and reduced to level 9 after 1 TR above the threshold, level 5 after 2 TRs above the threshold, and level 1 (minimal) after 3-5 TRs above the threshold. Participants also received a monetary reward and an emoji after the trial based on the final number of above-threshold TRs: 0 TRs means no money and an unhappy face; 1 TR means no money and a neutral face; 2-3 TRs means 5 cents bonus and a smiling face; and 4-5 TRs means 10 cents bonus and a laughing face.

Participants were informed about these types of feedback and that the feedback depended on their performance in the task. However, they were not instructed that the feedback was based on competitor activation, nor were they instructed on what mental strategy to use. Instead, they were instructed to explore different strategies that seemed to improve feedback. After the study, participants completed a debriefing questionnaire.

The threshold used for determining feedback was adjusted dynamically using an adaptive staircase procedure (Supplementary Table 1). The goal in using staircasing was to start participants at a difficulty level they could achieve at the beginning during their strategy exploration, but then to increase difficulty across runs and sessions such that they would be incentivized to activate the competitor object more and more strongly as they gained control of the feedback. When participants exhibited poor performance, the threshold was decreased, giving them an opportunity to improve and catch up. Conversely, the threshold was increased to create room for further improvement when participants demonstrated higher levels of control.

### Feedback ROI

The BOLD activity patterns used to train and test the object classifiers were extracted from a data-driven region of interest (ROI). To define this feedback ROI, we used the neuroSketch dataset [36], in which the same four objects were shown multiple times to other participants. Each of the 300 parcels in the Schaefer atlas [37] classified as gray matter by Freesurfer [38] were retained for further analysis. Individual classifiers were trained for each parcel, and their test performance was quantified using a leave-one-run-out methodology. The performance from each parcel was averaged across all 25 participants in the neuroSketch dataset, resulting in a ranking of parcel performance. To identify the set of parcels that yielded the best performance, we built a mega ROI adding in the voxels from the top-N parcels and re-calculating test performance for each value of N using a leave-one-run-out approach. The mega ROI composed of the 78 highest-performing parcels yielded the best object classification performance in the neuroSketch dataset.

This mega ROI with 78 parcels served as the starting point for each participant in the current study, but was further refined per individual through a greedy approach. We first removed the voxels from one parcel (77 parcels remaining) and trained and tested a 4-way classifier on the recognition runs from Session 1 with leave-one-run-out cross-validation, and then iterated until all 78 parcels had been the one parcel left out; the iteration in which the remaining 77 parcels yielded the highest decoding performance was retained. Then we repeated the whole procedure, dropping one parcel (76 parcels remaining), iterating until all 77 remaining parcels had been left out, and then retaining the best set of 76 parcels. This process repeated until the voxels from only one parcel remained, yielding performance values for mega ROIs containing 1–78 parcels; the mega ROI with the best performance of all of these combinations was used as the feedback ROI for this participant. As a result, different participants had a different number of parcels in their mega ROI. However, there was good consistency in which parcels were selected, especially in the visual cortex (Fig. 2a).

**Figure 2.**
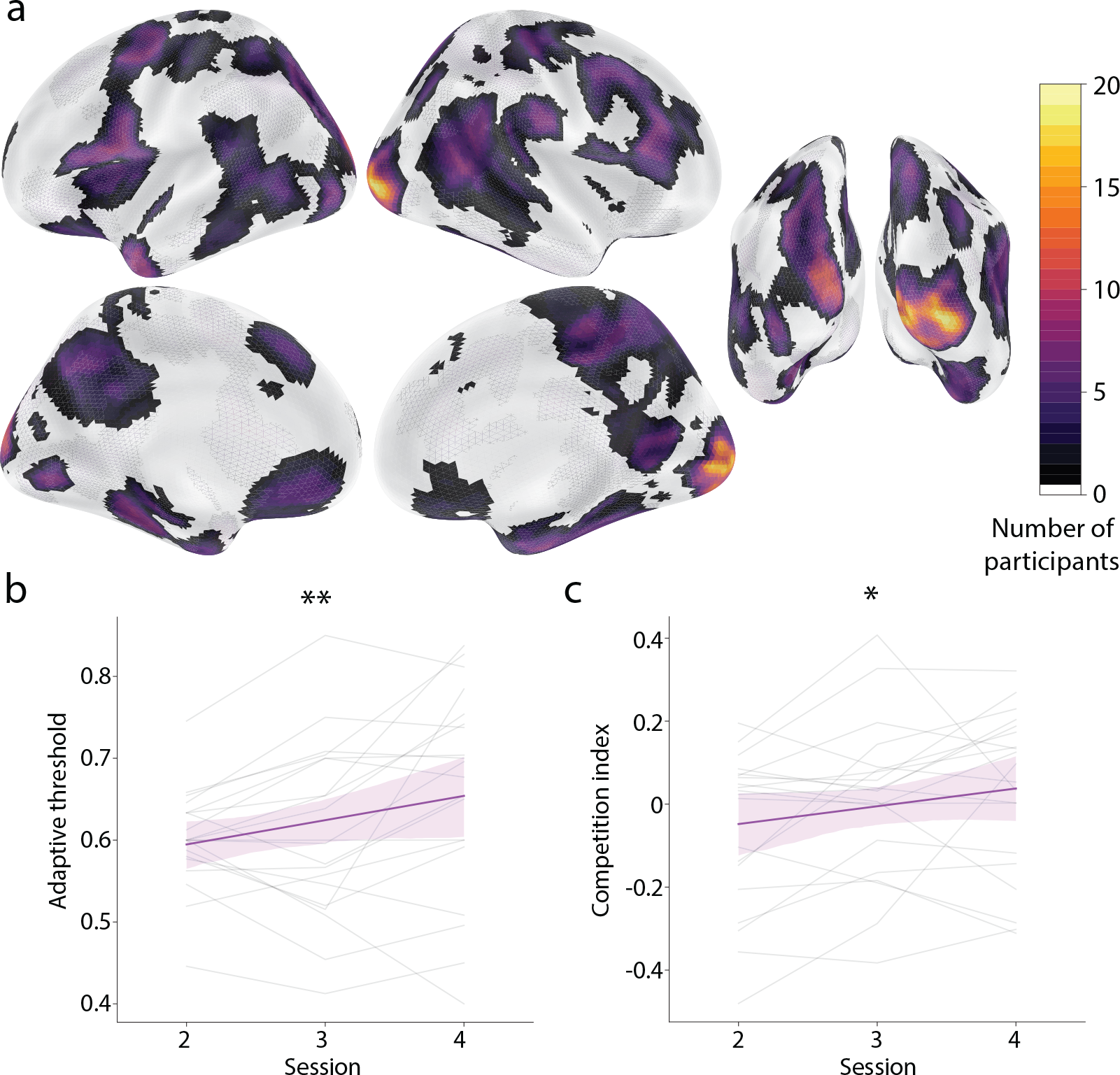
Neurofeedback validation. (a) The mega ROI used to provide neurofeedback was customized for each participant in a data-driven manner. This plot shows the number of participants whose feedback ROIs contained each voxel. The greatest consistency (virtually every participant) was observed in the early visual cortex, extending ventral and lateral. (b) Participants succeeded in self-generating classifier evidence for the competitor object based on neurofeedback, as indicated by an increase across the feedback sessions in the threshold from the staircasing procedure. A higher threshold over time indicates that participants were able to activate the competitor object more and more strongly. (c) As a corollary of this effect, we also found an increase across feedback sessions in the coactivation between the presented and competitor objects, quantified as the product of the classifier evidences for these objects relative to the control object baseline. *\*\** = *p <* 0.01, *\** = *p <* 0.05.

### Representational change ROIs

To examine how cortical coactivation related to representational change in the hippocampus, we segmented the hippocampus and its subfields anatomically for each participant using the automatic segmentation of hippocampal subfields (ASHS) software package [39] and a reference library of 51 manual segmentations [40, 41]. These segmentations defined participant-specific ROIs for the bilateral hippocampus and subfields CA1, CA2/3, dentate gyrus, and subiculum. We also explored broader effects in the cortex by using Freesurfer [42] to create masks for V1, V2, lateral occipital (LO) cortex, inferior temporal (IT) cortex, fusiform gyrus (FG) and parahippocampal cortex (PHC).

### Categorical perception task

To assess the impact of representational change on behavior, we conducted a categorical perception task [35, 29] before and after neurofeedback training (in Sessions 1 and 5, respectively). During this task, participants categorized images sampled from along a morph continuum between two object endpoints (e.g., bed to chair). Because the objects were rendered from 3-D models with a matching number of vertices, it was possible to smoothly morph between them. The morph percentage (of the second object) was sampled at 13 steps: 18 (i.e., 18% chair, 82% bed), 26, 34, 38, 42, 46, 50, 54, 58, 62, 66, 74, and 82. These morphs were shown 12 times each during both the pre-test and post-test, always from a trial-unique viewpoint. On each trial, participants were briefly presented (1 s) with the morph and asked to make a forced-choice judgment about which object they saw by clicking one of two buttons that appeared below the image. The assignment of labels to left vs. right buttons was randomized across trials. A logistic regression model was used to analyze the relationship between the morphing parameter and categorization responses:

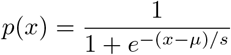

For the Session 1 categorical perception task, the slope and *μ* parameters were estimated. In Session 5, the *μ* value from Session 1 was fixed (to reduce noise and enhance model capability) and we estimated the slope. The change in slope indicates the type and degree of representational change between the two objects being discriminated. In particular, a decrease in slope (reduced discriminability) when comparing the presented and competitor objects would be consistent with integration of their representations, whereas an increase in slope (improved discriminability) would be consistent with differentiation. We thus defined a behavioral integration index as the Session 1 slope minus Session 5 slope (positive for integration, negative for differentiation). Importantly, these changes can be evaluated with respect to the analogous change observed for the untrained control objects.

### Representational change analysis

We examined representational change in the hippocampus and elsewhere by comparing the overlap of neural patterns for the presented and competitor objects (using control objects 1 and 2 as a baseline) in Session 1 vs. Session 5. For Session 1, we built eight regularized logistic regression classifiers, each using seven of the recognition runs for training and the final recognition run for testing in a leave-one-run-out manner. The test accuracies of the eight classifiers were averaged to compute an overall score. For Session 5, we tested the eight trained classifiers from Session 1 on the eight recognition runs and again averaged their accuracies to compute an overall score. Higher classification accuracy indicates less neural overlap, and thus a decrease in accuracy from Session 1 to Session 5 is consistent with integration and an increase in accuracy is consistent with differentiation. We thus defined a neural integration index for each ROI as the Session 1 accuracy minus Session 5 accuracy (positive for integration, negative for differentiation).

### Brain-behavior relationship

To the extent that categorical perception is a behavioral readout of neural overlap in one or more ROIs, the behavioral and neural integration scores should be positively correlated across participants. We quantified this association in each ROI by calculating the Pearson correlation coefficient.

### Statistics

We used non-parametric statistics where possible to avoid assumptions of parametric tests. To estimate the sampling distribution of an effect, we performed bootstrap resampling at the group level. Namely, from the original sample of 20 participants, for each of 1,000 iterations we sampled 20 participants with replacement and averaged their values. The mean and 95% bounds of the resulting sampling distribution were used to generate the bar plot. For hypothesis testing, we determined the p-value as the proportion of iterations on which the average had the opposite sign from the original effect.

## Data availability

Data and analysis code will be shared publicly upon acceptance.

## Results

### Validation of real-time neurofeedback

We sought to use real-time fMRI neurofeedback to generate coactivation between an object being perceived (presented object) and an object being internally activated from memory (competitor object). We tested whether participants succeeded in using neurofeedback to activate the competitor object in the feedback sessions (Sessions 2–4) in two ways. First, we examined how the adaptive threshold from the feedback staircasing procedure changed over time; an increase in the threshold would indicate that participants were able to activate the competitor object more and more strongly. Indeed, using linear regression we found a reliable increase in the adaptive threshold across feedback sessions (Fig. 2b, *p* = 0.004). Second, we more directly tested whether coactivation between the presented and competitor objects increased over time. We quantified coactivation by multiplying the classifier evidence from the feedback ROI for the presented object (relative to control objects) times the classifier evidence for the competitor object. Again, we found a significant rise in this neural “competition index” over feedback sessions (Fig. 2c, *p* = 0.025).

### Behavioral integration

We next examined the impact of this induced coactivation on the representations of the presented and competitor objects from before (Session 1) to after (Session 5) neurofeedback training. We used a categorical perception task [35] as a behavioral assay of the overlap between representations. Participants were shown a morph of two objects and had to indicate which of the two objects they saw by selecting the label (Fig. 3a). To the extent that coactivation led to integration, greater reliance on the shared features of the two objects should make it harder for participants to discriminate the morph and lead to a shallower logistic slope; alternatively, if differentiation occurred, then greater reliance on the unique features of the two objects should exaggerate those features when perceiving the morph and lead to easier discrimination and a steeper slope.

**Figure 3.**
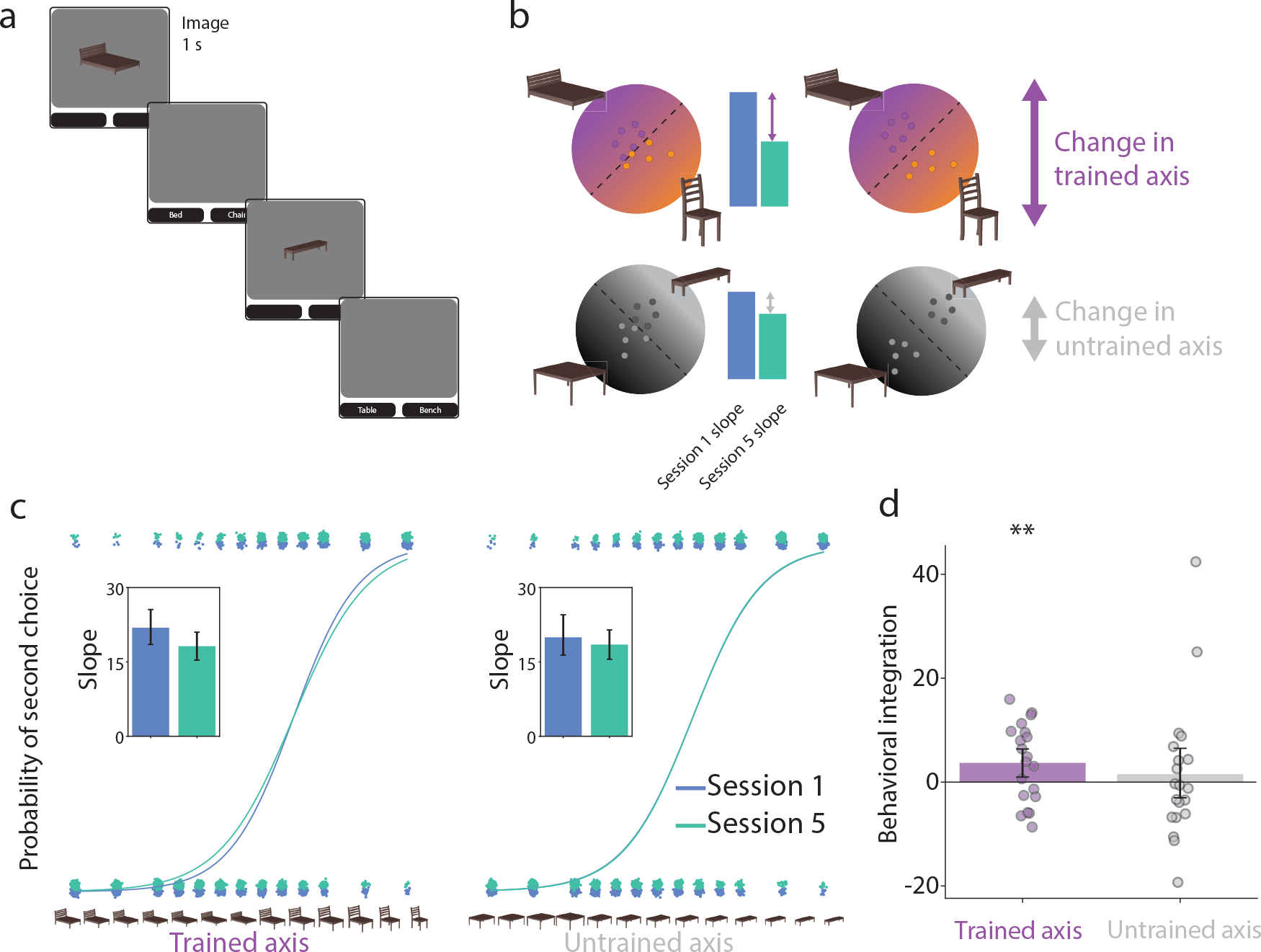
Categorical perception. (a) On each trial of the categorical perception task, participants were shown a morph on a continuum between two objects and asked to indicate via forced choice responses which of the two objects they perceived. (b) Behavioral categorization responses can be modeled with logistic regression as a function of morph level. For each participant, separate models were fit to the first (Session 1) and last (Session 5) session and to discrimination of trained objects (presented vs. competitor) and of control objects (control 1 vs. 2). (c) The fitted slope parameters (see insets) were then used to calculate a behavioral integration index of Session 1 minus Session 5. (d) This index was reliably positive for the trained axis, reflecting a shallower slope after neurofeedback indicative of integration. However, there was no reliable change for the untrained axis, suggesting that neurofeedback was necessary for integration. Each dot represents an individual participant and the error bands reflect the 95% confidence interval from bootstrap resampling. *\*\** = *p <* 0.01.

A logistic regression was fitted to the categorization responses as a function of the morphing parameter (Fig. 3c), separately for Sessions 1 and 5, and for the trained (presented vs. competitor objects; e.g., bed, chair) and untrained (control 1 vs. 2 objects; e.g., bench, table) axes. We calculated a behavioral integration index as the difference in the slope for Session 1 minus Session 5 (Fig. 3b); positive indicates that the slope got shallower. This index was reliably positive for the trained axis (*p* = 0.009; Fig. 3d), but not for the untrained axis (*p* = 0.314); the difference in indices between trained and untrained axes did not reach significance (*p* = 0.242). This provides behavioral evidence of integration between the presented and competitor objects as a result of neurofeedback.

### Neural integration

We hypothesized the coactivation of object representations in the cortical inputs to the hippocampus would lead to representational changes in the hippocampus. According to the non-monotonic plasticity hypothesis (NMPH), strong coactivation should lead to integration, whereas moderate coactivation should lead to differentiation. To examine these representational changes, we applied binary classifiers to patterns of BOLD activity from hippocampal ROIs before (Session 1) and after (Session 5) neurofeedback training using cross-validation (Fig. 4a). If two objects integrate, their greater neural overlap should lead to misclassifications and reduced accuracy; in contrast, differentiation and lower overlap should make classification easier and increase accuracy. We computed classification accuracy in the recognition runs from Sessions 1 and 5 separately for the trained axis (presented vs. competitor) and untrained axis (control 1 vs. 2). As with behavioral integration, we calculated a neural integration index for each ROI as the classification accuracy in Session 1 minus Session 5 (positive indicates classification worsened).

**Figure 4.**
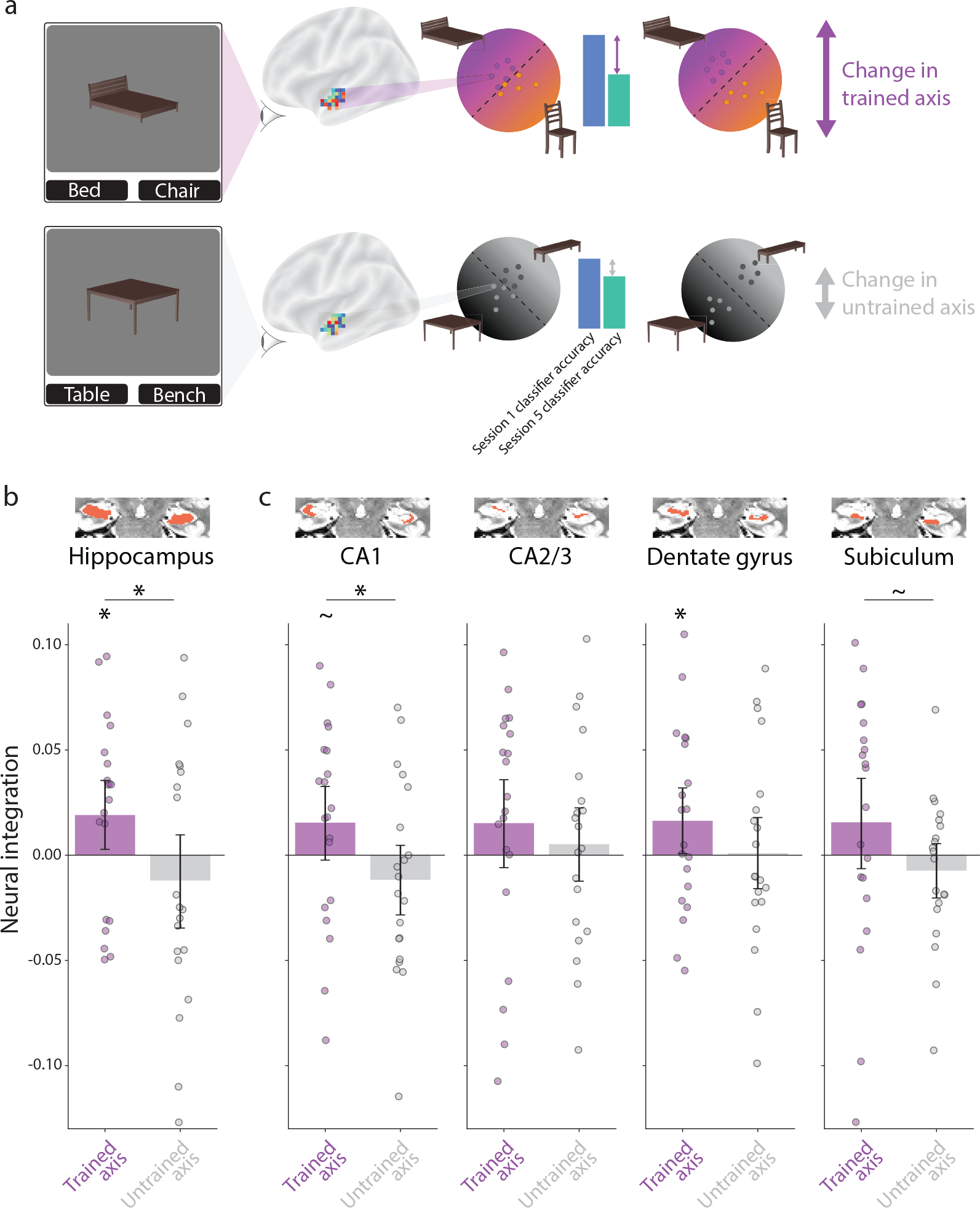
Representational change in the hippocampus. (a) We collected neural snapshots of how the four objects were represented in each ROI before (Session 1) and after (Session 5) neurofeedback training. Participants viewed the objects (intact, no wobbling or morphing) during a recognition task in these sessions. For each participant, we built regularized logistic regression classifiers to discriminate multivoxel patterns of BOLD activity in each ROI evoked by the presented vs. competitor objects (trained axis) and by the control 1 vs. 2 objects (untrained axis). Using leave-one-run-out cross-validation, we quantified the neural overlap along these two axes from the test classifier accuracy (high accuracy indicates less overlap). A neural integration index was defined for each ROI and axis as classifier accuracy in Session 1 minus Session 5 (positive reflects a decrease in classifier accuracy). (b) In the whole hippocampus, there was reliable neural integration along the trained axis relative to the untrained axis. The integration for the trained but not untrained axis was reliably above chance. (c) The difference in neural integration between trained and untrained axes was reflected most clearly in the CA1 subfield, though also marginally in the subiculum, and the trained axis was above chance in the dentate gyrus. The insets depict the location of the ROI in red on an example participant’s T2 scan. Each dot in the bar plots represents an individual participant and the error bands reflect the 95% confidence interval from bootstrap resampling. *\** = *p <* 0.05.

In the overall hippocampus (Fig. 4b), there was a significant difference between indices for the trained and untrained axes (*p* = 0.043); the trained axis (*p* = 0.027) but not the untrained axis (*p* = 0.819) was reliably positive. These results indicate that integration occurred as a result of neurofeedback in the hippocampus. Looking at individual subfields, we were especially interested in the CA1 subfield, which has previously been linked to integration [21, 11]. Indeed, as in the overall hippocampus, CA1 was the only subfield to show a reliable difference in neural integration between trained and untrained axes (*p* = 0.016); subiculum was marginal (*p* = 0.063), none of the other subfields reached significance (*ps >* 0.132). The trained axis on its own was marginal (*p* = 0.075) in CA1 though reached significance in the dentate gyrus (*p* = 0.042); the remaining subfields and the untrained axis in all subfields did not differ from chance (*ps >* 0.117).

In the cortical ROIs, the PHC mirrored the hippocampus in showing integration as a result of neurofeedback, with a significant difference in neural integration index between trained and untrained axes (*p* = 0.009), a reliably positive index for the trained axis (*p* = 0.009), and no difference from chance for the untrained axis (*p* = 0.874). None of the other cortical ROIs showed a difference in neural integration indices between trained and untrained axes (*ps >* 0.115).

### Relationship between behavioral and neural integration

Given that we observed both behavioral and neural integration after neurofeedback training, we next asked if these effects are related (Fig. 5). Namely, does the increased neural overlap of the trained objects in the hippocampus, CA1, and PHC have behavioral significance for categorical perception? For the trained axis, we found a marginal positive correlation between neural integration in the hippocampus and behavioral integration (*r* = 0.383, *p* = 0.095). This brain-behavior relationship was a bit clearer in PHC (*r* = 0.445, *p* = 0.049), but not found in CA1 (*r* = 0.020, *p* = 0.936). As a control, we repeated this analysis for the untrained axis and found no reliable correlations (*ps >* 0.403).

**Figure 5.**
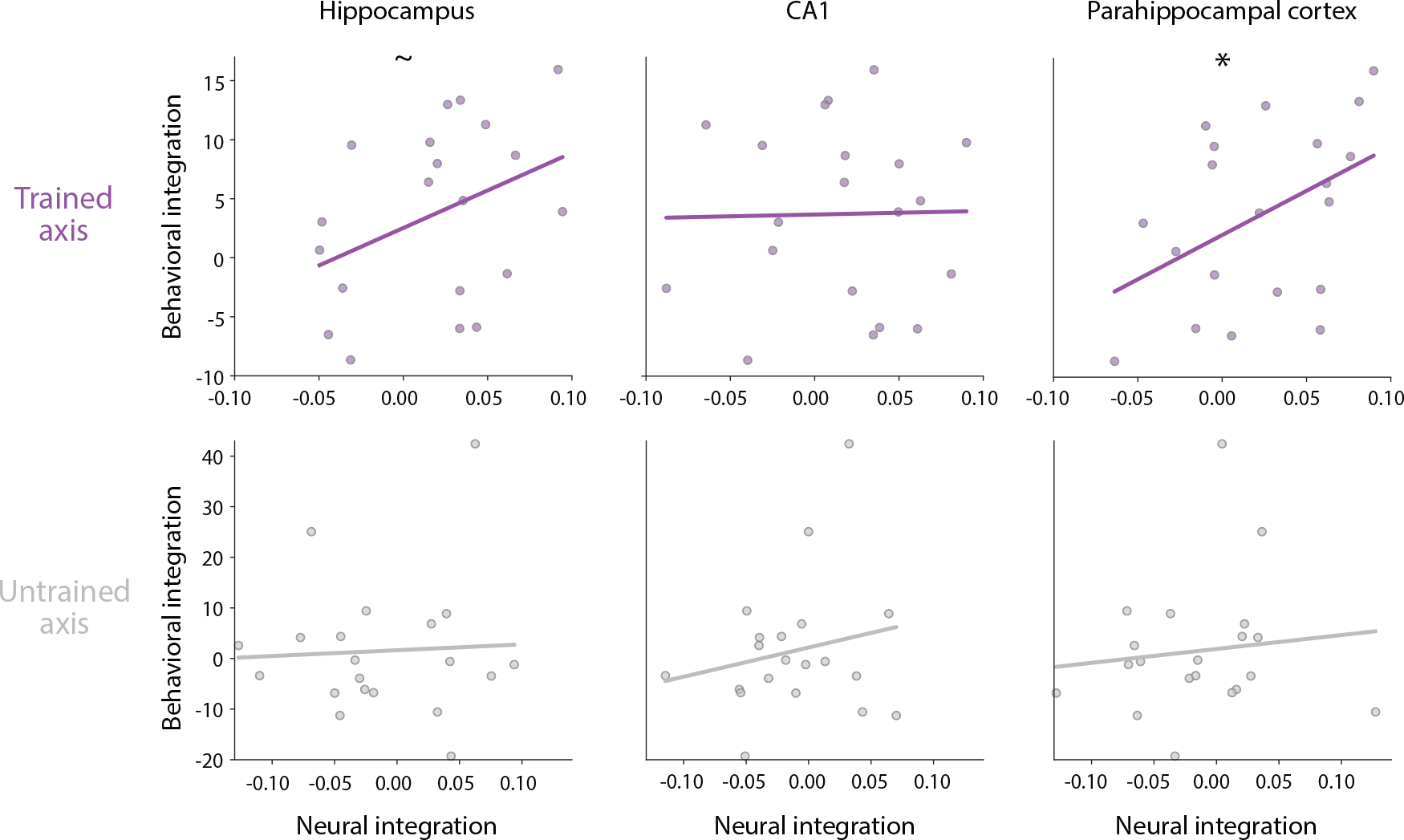
Behavioral relevance of neural representational change. The amount of neural integration predicted the amount of behavioral integration in PHC and marginally in the hippocampus, though not reliably in CA1. Each dot represents an individual participant and the line indicates the fitted linear relationship. *\** = *p <* 0.05, = *p <* 0.10.

## Discussion

The current study used closed-loop neurofeedback from real-time fMRI to provide a direct test of how the coactivation of object representations in the visual cortex drives representational change in the hippocampus. The key findings are that: (1) Participants succeeded in using implicit neurofeedback to increase activation of one representation (the competitor object) while viewing another object (the presented object) across three training sessions; (2) this intervention for one pair of objects (trained axis) led participants to be more easily confused in a categorical perception task administered in pre- and post-training sessions, which was not observed for a control pair of objects (untrained axis); (3) neural snapshots of the representations of the trained object pairs in the hippocampus, CA1, and PHC were harder to classify in the post-vs. pre-training sessions; and (4) the amount of this neural integration in PHC (and marginally in hippocampus) was correlated across participants with reduced behavioral discriminability in the categorical perception task. These results indicate that increasing coactivation of objects in the visual cortex can drive their neural representations in the medial temporal lobe to integrate and that this can have behavioral consequences for perception.

We interpret the observed neural integration in the hippocampus as consistent with the right tail of the nonmonotonic plasticity hypothesis [3]. That is, by encouraging participants to activate the competitor objects more and more strongly with a staircasing procedure, and through task and monetary incentives, the activation levels of the presented and competitor objects may have been high enough to foster synaptic strengthening between their shared features and unique features of each object, allowing viewing of either object to reactivate the hippocampal representation of the other object. This in turn would increase the similarity of the patterns of fMRI activity evoked by the individual objects during the recognition runs.

The NMPH also predicts that moderate coactivation would lead to the opposite effect of differentiation, via synaptic weakening between shared and unique features. A different feedback regime in a future study may be more effective at targeting moderate coactivation. Namely, rather than trying to maximize competitor activation, the amount of activation could be titrated with neurofeedback to keep it from getting too high (or too low). Different groups of participants, or even different object pairs within participants, could have different low-medium-high target activation levels to span the NMPH curve more continuously. Because hippocampal subfields and other brain regions differ in activation profiles, this may lead to more varied integration and differentiation results. Finally, for simplicity we provided feedback based on competitor activation, but the hypothesis requires coactivation. We assumed that presenting an object perceptually would be sufficient to ensure coactivation with the competitor (as shown in Fig. 2b), but future studies could provide neurofeedback on coactivation rather than just competitor activation.

These findings that internally generated competition can prompt neural integration in the hippocampus provide key new evidence for theories of memory. Previous studies of representational change in the hippocampus [14, 18, 15, 13, 11] found differentiation and/or integration results consistent with the NMPH. In these studies, coactivation was achieved through a stimulus or task manipulation, such as by increasing stimulus similarity [14] or by encouraging retrieval of related experiences [11]. Here we directly targeted coactivation by using neurofeedback to incept activation of a competing representation. Following the logic of prior real-time fMRI studies [28], this demonstrates that neural coactivation itself is sufficient for hippocampal integration. This is a stronger and more causal test of the hypothesis that neural competition drives representational change.

The role of the hippocampus, and CA1 in particular, in memory integration accords with existing findings [11] and computational models [21]. CA1, which receives cortical input from the entorhinal cortex along the monosynaptic or temporoammonic pathway, is a key site for integration because it has lower inhibition between neurons, which allows more neurons to become active and increases baseline overlap across memories. As a result, CA1 is more likely to enter a regime in which a target and competitor memory are strongly active, strengthen their synaptic interconnections, and produce more integrated memories. PHC, a key relay between the visual cortex and the hippocampus, has also previously been implicated in memory integration [43]. However, failures of integration in patients with hippocampal lesions (some with intact parahippocampal cortex) raise the question of whether PHC is sufficient for behavioral integration [44], despite it showing the strongest the brain-behavior correlation in the current study. We also observed a marginal neural integration effect in the subiculum, a structure traditionally considered a relay from the hippocampus to other brain regions[45]. Indeed, the subiculum is a crucial link to the brain’s reward system, which was likely engaged by the neurofeedback received[46]. An exciting direction for future research could be to examine how reward processing, for example in the ventral tegmental area (VTA)[47] and ventral striatum[48, 49, 50], can modulate coactivation in visual cortex and representational change in the hippocampus.

In sum, this study employs real-time neurofeedback to induce memory competition and drive representational changes in the hippocampus. This approach may be more efficient than using predefined stimuli or tasks to manipulate competition, as it personalizes neurofeedback to the most proximal variable of interest — cortical activation of the competing memory. With further development and improved neural technologies, this technique could find a variety of applications. For example, it could strengthen memories in age-related memory loss or weaken memories in PTSD by reducing or increasing competition, or it could be used to combine or disambiguate representations of related concepts in stroke patients or even in educational contexts.

## Acknowledgements

We thank Wenyan Bi, Aalap Shap, Qi Lin, Tristan Yates, Hillary Nguyen, Sheri Choi, Ariadne Letrou, Xihan Zhang, Erica Busch, Brynn Sherman, Kathryn Graves, and Ben Swinchoski for their help with data collection.

---

**Supplementary Table 1.** Staircase procedure for adaptive threshold. Trials were considered successful when 2 or more of the TRs (of 5 total feedback TRs per trial) were above the threshold.

| procedure |  |  |  |  |  |
| --- | --- | --- | --- | --- | --- |
| if first neurofeedback training run of session 1 |  |  |  | then | threshold=60% |
| else if first neurofeedback training run of day N |  |  |  |  | threshold = last threshold from day N-1 |
| else if<br>number of<br>successful<br>trials | <=1 | for the previous | 1 run |  | decrease threshold by 5% (min 40%) |
|  | <=3 |  | 3 runs |  |  |
|  | <=5 |  | 5 runs |  |  |
|  | ==6 |  | any runs |  | keep threshold unchanged |
|  | >=7 |  | 5 runs |  | increase threshold by 5% (max 90%) |
|  | >=9 |  | 3 runs |  |  |
|  | >=11 |  | 1 run |  |  |

